# Intermittent Fasting Reprograms Chromatin Accessibility to Modulate Gene Expression in Brain and Muscle

**DOI:** 10.1101/2025.02.11.637778

**Authors:** Yibo Fan, Xiangyuan Peng, Mansour Ebrahimi, Nishat I. Tabassum, Xiangru Cheng, Yatendra Kumar, Yong U. Liu, Guobing Chen, Eitan Okun, Terrance G. Johns, Christopher G. Sobey, Mitchell K. P. Lai, Jayantha Gunaratne, Mark P. Mattson, Dong-Gyu Jo, Thiruma V. Arumugam

## Abstract

Intermittent fasting (IF), a dietary regimen that mimics the natural feeding patterns observed across diverse organisms—from single-celled life to mammals—is thought to activate systemic survival responses and confer health benefits. However, the molecular mechanisms underlying these effects remain poorly understood. In this study, we utilized ATAC-Seq and RNA-Seq to investigate how IF influences chromatin accessibility and gene expression in brain and muscle tissues of mice, compared to ad libitum feeding. Our results reveal that IF induces significant changes in chromatin accessibility, modulating pathways related to metabolism, ribosome function, HIF-1 signaling, and glycolysis. Motif analysis identified tissue-specific transcription factors enriched in IF-regulated regions, including Sp1, Mef2a, NeuroD2, Banp, and NFIA in the brain, and SMAD4, TCF4, STAT5B, NKX3-1, and ZEB2 in muscle. Integrative analysis of ATAC-Seq and RNA-Seq data demonstrated that IF upregulates 50 genes and downregulates 15 in the cortex, while upregulating 31 genes and downregulating 10 in muscle. These gene expression changes are linked to pathways associated with neuroprotection and enhanced muscle function, offering mechanistic insights into the health benefits of IF. Our findings underscore the role of IF-induced chromatin remodeling in driving adaptive gene regulation.

## Introduction

Intermittent metabolic switching, induced by interventions such as exercise, caloric restriction, and intermittent fasting (IF) (e.g., time-restricted feeding), has been demonstrated to confer significant health benefits across various species. These metabolic shifts are associated with an extension of both healthspan and lifespan. IF typically involves periods of 12 to 24 hours with minimal or no caloric intake, resulting in a metabolic transition from glucose derived from hepatic sources to the utilization of fatty acids and ketones (1). Over the past few decades, extensive animal research has established that IF protects against metabolic disorders (2), age-related neurodegenerative diseases (3-6), and cardiovascular conditions (7). Additionally, IF has been implicated in offering protective effects against specific types of cancer (8). Notably, pioneering studies conducted by our group and collaborators have demonstrated that every-other-day (EOD) fasting and 16-hour time-restricted fasting are effective in mitigating Alzheimer’s disease (AD) (6), vascular dementia (4, 9), heart failure (7), and stroke (3, 5, 10-12). Recently, an increasing number of studies have begun to investigate the effects of IF in human populations, contributing to a deeper understanding of its potential therapeutic benefits (13-15). In a recent 8-week randomized clinical trial involving 40 cognitively intact older adults with insulin resistance, IF was found to improve biomarkers associated with insulin signaling, as well as blood markers of carbohydrate and lipid metabolism (16). Additionally, IF positively impacted executive function and memory (16). Studies on skeletal muscle suggest that IF activates signaling pathways that enhance cellular stress resistance, mitochondrial biogenesis, and autophagy (17).

As the beneficial effects of IF on health and lifespan have been well-documented, we employed multi-omics approaches to investigate the molecular mechanisms underlying the metabolic switch induced by IF. RNA sequencing studies in mice yielded comprehensive data on how IF influences the expression of genes that remodel key organs, including the liver, heart, and brain (10, 18-20). Furthermore, these studies provided insights into the protective mechanisms of IF against ischemic stroke (10), revealing that IF differentially regulates genes involved in cell survival, inflammation, neuroplasticity, and neurogenesis to confer protection against stroke (10). IF-induced changes in gene expression may be controlled by epigenetic mechanisms. Studies investigating epigenetic regulation have revealed that IF can influence histone modifications and the DNA methylation landscape, thereby regulating gene expression (19). Specifically, IF has been shown to modulate trimethylation of histone H3 at lysine 9 (H3K9me3) in the brain, orchestrating a wide array of transcriptomic changes associated with robust metabolic switching processes (19). Notably, both the epigenomic and transcriptomic alterations induced by IF were retained following refeeding, suggesting that the memory of IF-induced epigenetic changes persists (19). Additionally, IF significantly increased the abundance of DNA methyltransferases (DNMTs), which could suggest enhanced hypermethylation through the addition of methyl groups (4). However, the overall DNA methylation pattern in IF-treated animals exhibited a distinct shift toward a hypomethylated state (4). Recent studies in mice have further explored whether changes in gene expression also translate into alterations in the proteomic and phosphoproteomic landscapes in response to IF (20). These investigations revealed that IF significantly impacts pathways involved in cyclic GMP signaling, lipid and amino acid metabolism, cell adhesion, apoptosis, and inflammation (20). However, the mechanisms by which IF regulates chromatin accessibility to influence transcriptional programs remain unknown.

Transcriptional programs are closely linked to alterations in chromatin states, which encompass a variety of epigenetic modifications to DNA, localized changes in chromatin structure, shifts in the three-dimensional topology of chromatin domains, and the overall organization of chromosomes within the cell nucleus (21-22). Among these features, local chromatin accessibility has emerged as a critical indicator of structural dynamics. Assay for Transposase-Accessible Chromatin using Sequencing (ATAC-seq) has rapidly established itself as a powerful technique for profiling chromatin accessibility. This innovative method employs hyperactive Tn5 transposase to simultaneously cleave DNA and insert sequencing adapters, targeting regions of open chromatin for analysis (23). In this study, we explored for the first time the impact of IF on chromatin accessibility in the brain and muscle in mice and examined the interconnections between IF-induced gene expression and chromatin accessibility in these two organs. Our results demonstrate that 16-hour daily IF over a 4-month period significantly alters chromatin accessibility and modifies pathways related to metabolism, ribosome function, HIF-1 signaling, and glycolysis in both the brain and muscle. Through comprehensive motif enrichment analysis, we identified key regulatory mechanisms underlying the effects of IF. Furthermore, integrative analysis of chromatin accessibility peaks and differentially expressed genes confirmed the observed changes in chromatin structure across both tissues. These findings provide valuable insights into the interplay between chromatin accessibility and gene expression changes that may contribute to the health benefits of IF in brain and muscle. This research advances our mechanistic understanding of IF and highlights its potential for translation into therapeutic strategies to promote health and combat age-related diseases.

## Materials and Methods

### Animals and IF Procedures

All *in vivo* experimental procedures were approved by La Trobe University (Ethics approval number: AEC21047) Animal Care and Use Committee and conducted in accordance with the guidelines outlined in the Australian Code for the Care and Use of Animals for Scientific Purposes (8th edition) and the NIH Guide for the Care and Use of Laboratory Animals. Every effort was made to minimize suffering and reduce the number of animals used. All sections of the manuscript were performed in accordance with ARRIVE guidelines (24). C57BL/6 male mice were purchased at 4 weeks of age from Animal Resources Centre (ARC), Australia, and housed in the animal facility at La Trobe University. The animals were subjected to a 12-hour light:12-hour dark cycle (07:00-19:00) and provided with normal diet comprising 20% total crude protein, 5% crude fat, 6% crude fiber, 0.5% added salt, 0.8% calcium and 0.45% phosphorus (Ridley). Water was available *ad libitum* to all dietary groups. At 6 weeks of age, male mice were randomly assigned to different dietary intervention groups: Intermittent Fasting for 16 hours (IF) and *ad libitum* feeding as a control (AL). Mice in the IF group underwent daily fasting for 16 hours for 4 months (16:00-08:00), while the AL group had continuous access to food pellets. After 4 months of dietary intervention, all mice were euthanized by carbon dioxide (CO2) inhalation between 7 a.m. and noon. Subsequently, all mice were perfused with cold PBS, and tissue samples were collected and snap-frozen in liquid nitrogen and stored at -80 °C until further use.

### Transposase-Accessible Chromatin Assay with Sequencing (ATAC-Seq)

ATAC-seq was performed using the Active Motif ATAC-seq kit (USA) according to the manufacturer’s instructions for total DNA extraction from muscle and cortex samples. Nuclei were isolated from the samples at a concentration of 50,000 to 100,000 cells. The nuclei pellet was then subjected to tagmentation and purification using the Tagmentation Master Mix. For library amplification, we employed four indexed I7 primers and I5 primers in a PCR reaction. Following PCR, libraries were purified using solid-phase reversible immobilization (SPRI) beads, and their quality was assessed using D1000 ScreenTape. The initial quantification of the libraries was performed on a TapeStation 4150, diluting the libraries to a concentration of 2-4 ng/μL. Next, we determined the insertion size of the libraries using NGS3K and quantified their accurate concentration through Q-PCR, ensuring that the effective concentration was greater than 2 nM for optimal pooling prior to sequencing. After completing quality control for the libraries, sequencing was conducted on different libraries based on their concentration and data requirements, utilizing the Illumina NovaSeq 6000 platform. Clustering of the index-coded samples was carried out on a cBot Cluster Generation System with the TruSeq PE Cluster Kit v3-cBot-HS (Illumina), following the manufacturer’s guidelines. After cluster generation, the library preparations were sequenced on the Illumina HiSeq platform, generating 150 bp paired-end reads.

### ATAC-Seq Data Analysis

Nextera adapter sequences were first trimmed from the reads using Skewer (v0.2.2). The quality of the raw reads was assessed with FastQC (v0.11.5). The trimmed reads were aligned to a reference genome using BWA (v0.7.12-r1039) with standard parameters. High-quality reads were filtered based on the following criteria: a mapping quality score (MAPQ) of ≥ 13, non-mitochondrial chromosomes, and properly paired reads longer than 18 nt. To analyze the ATAC-seq signal, deepTools (v3.0.2) was employed to evaluate the regions 3 kb upstream and downstream of the transcription start site (TSS), as well as at peak summits and within gene bodies. Peak calling was conducted using MACS2 (v2.1.2) with a threshold q-value of 0.05. The distribution of peaks across various functional areas was analyzed using the ChIPseeker software. To investigate sequence information, HOMER (v4.9.1) was utilized to identify conserved sequence features by analyzing 250 bp (totaling 500 bp) upstream and downstream of the peak regions. The most significant motifs of lengths 8, 10, 12, and 14 were identified for both the IF and AL groups. Finally, the Integrative Genomics Viewer (IGV, v2.18.0) was used to visualize the alignment of corresponding transcription factors of the most enriched motifs.

### Generation of Heat Maps and Functional Enrichment Analysis

R and the pheatmap package were utilized, along with the fold enrichment output from MACS2 (v2.1.2), to generate heatmaps of differentially expressed peaks. To assess potential biological functional changes arising from the observed gene expression alterations, we performed Gene Ontology (GO) enrichment analysis using the GOseq and topGO R package (v4.10.2), which corrects for gene length bias. For KEGG pathway enrichment analysis, we employed the KEGG Orthology Based Annotation System (KOBAS 3.0) software to evaluate statistical enrichment. GO terms and KEGG pathways with adjusted p-values less than 0.05 were deemed significantly enriched among the DEGs.

### Ribonucleic Acid (RNA) Extraction

Total RNA was extracted from muscle and cortex samples using the TRIzol RNA Isolation Kit (Thermo Fisher Scientific, USA), following the manufacturer’s instructions. The quality of the extracted RNA was assessed by measuring RNA purity with a Nanodrop ND-1000 spectrophotometer (Thermo Fisher Scientific, USA). High-quality RNA was indicated by an OD260/OD280 ratio of 1.9–2.0, as determined from the Nanodrop readings.

### cDNA Library Construction and RNA Sequencing (RNA-Seq)

The NEBNext Ultra II RNA Library Prep Kit for Illumina and the NEBNext® Ultra™ II Directional RNA Library Prep Kit (New England BioLabs, USA) were utilized following the manufacturer’s instructions for eukaryotic RNA sequencing (RNA-seq). Poly-T oligo-attached magnetic beads were employed to purify mRNA from total RNA samples. Following purification, a fragmentation buffer was added to the mRNA. First-strand complementary DNA (cDNA) synthesis was carried out using random hexamer primers. Subsequently, second-strand complementary RNA (cRNA) synthesis was performed using deoxyuridine triphosphate (dUTP) to facilitate directional library preparation. After the completion of end repair, A-tailing, adapter ligation, size selection, USER enzyme digestion, amplification, and purification, the library preparation yielded a minimum of 12 GB of raw data per sample (Illumina, USA). The quality of the library was assessed using Qubit for quantification and real-time PCR for verification, alongside Bioanalyzer analysis for size distribution profiling. Quantified libraries were pooled and sequenced on Illumina platforms, based on effective library concentration and required data volume.

### Transcriptome Data Mapping and Analysis of Differential Expression

The RNA sequencing results generated by the Illumina NovaSeq 6000 Sequencing System were output as quality and color space FASTA files. These files were mapped to the Ensembl-released mouse genome sequence and annotation (Mus musculus; GRCm39/mm39). The HISAT2 v2.0.5 software was employed to build an index of the reference genome and to align the paired-end clean reads to it. Gene expression levels were quantified using FeatureCounts v1.5.0-p3, which counted the number of reads mapped to each gene. Fragments per kilobase of transcript per million mapped reads (FPKM) for each gene were then calculated based on the length of the gene and the corresponding read counts. Differential expression analysis was conducted using the DESeq2 R package (v1.10.1). Genes with a p-value < 0.05, as identified by DESeq2, were classified as differentially expressed.

### Functional Enrichment Analysis

To assess the potential biological functional changes resulting from gene expression alterations, Gene Ontology (GO) and Kyoto Encyclopedia of Genes and Genomes (KEGG) pathway enrichment analyses were conducted. Both GO and KEGG enrichment analyses were performed using the clusterProfiler R package (version 3.8.1), with corrections made for gene length bias. GO terms and KEGG pathways with an adjusted p-value of less than 0.05 were considered significantly enriched among the differentially expressed genes (DEGs).

### Integrative Analysis of Chromosome Accessibility and Transcriptome Data

For the integration of chromosome accessibility and transcriptome data, only differentially expressed genes (DEGs) with a p-value of less than 0.05, along with corresponding aggregated chromosome accessibility peaks (|log2 foldchange| ≥1), were included in the analysis. To compare changes in transcriptome and chromosome accessibility between the IF and AL groups, the VennDiagram R package (version 3.0.3) was utilized.

### Heat Map Generation Analyses of Integrative Data

R and VennDiagram package (v1.7.3) were used to filter common and differentially expressed genes from the differentially expressed peaks from MACS2 (v2.1.2) in ATAC-seq and DEGs from DESeq2 (v1.20.0) from RNA-seq. R and the pheatmap package (v1.0.12) were utilized for heatmap of common expressed genes.

### Statistical Analysis

For physiological measurements such as body weights and bodyweight change, values are indicated as mean ± SEM (n=8 mice/group). Significance is based on two-way ANOVA with Tukey’s post-hoc test. P-value <0.05 was considered statistically significant. For physiological measurements, such as fasting blood glucose, ketone levels, values are indicated as mean ± SEM (n=8 mice/group). Significance is based on two-tailed paired T-test. P-value <0.05 was considered statistically significant. The statistics for these comparisons were conducted using the GraphPad Prism software programme. For chromatin accessibility data, MACS2 software (threshold q value = 0.05) is used to call peak. |Log2foldchange| >1 is defined as differential expressed peaks. Differential analysis is performed using FoldEnrich values of different group peaks. Differential binding sites are analyzed in different groups by finding differential peaks when the ratio of FoldEnrich is larger than 2. For these significant peaks related genes, functional enrichment based on GO terms and associated KEGG pathways were inferred based on adjusted p-values <0.05 as implemented in Goseq, topGO v4.10.2 and KOBAS v3.0. For transcriptome data, differential expression analysis of the quantified transcripts was performed using DESeq2 R Package v.1.20.1. Those transcripts with p-value <0.05 were considered to be significantly differentially expressed among the different categories. For these significant genes, functional enrichment based on GO terms and associated KEGG pathways were inferred based on adjusted p-values <0.05 as implemented in clusterProfiler v3.8.1.

## Results

The study design, including the timing of experimental interventions and blood and tissue collections, is summarized in **Figure 1a**. Male C57BL/6 mice were fed a normal diet comprising 20% total crude protein, 5% crude fat, 6% crude fiber, 0.5% added salt, 0.8% calcium, and 0.45% phosphorus. Mice were randomly assigned to AL or daily IF schedules beginning at 6 weeks of age. IF animals exhibited significantly lower body weight and body weight changes compared to AL mice during the four-month dietary intervention period (**Figure 1b-c**). To assess the impact of IF on energy metabolism, blood glucose, ketone levels, HbA1c and cholesterol were measured in all mice. Mice in the AL group had significantly increased blood glucose and HbA1c levels, whereas IF mice had significantly decreased blood glucose levels (**Figure 1d-e**). Blood ketone levels were significantly higher in the IF group compared to the AL group (**Figure 1f**). Additionally, IF mice had significantly lower cholesterol levels compared to AL mice (**Figure 1g**).

**Figure 1:**
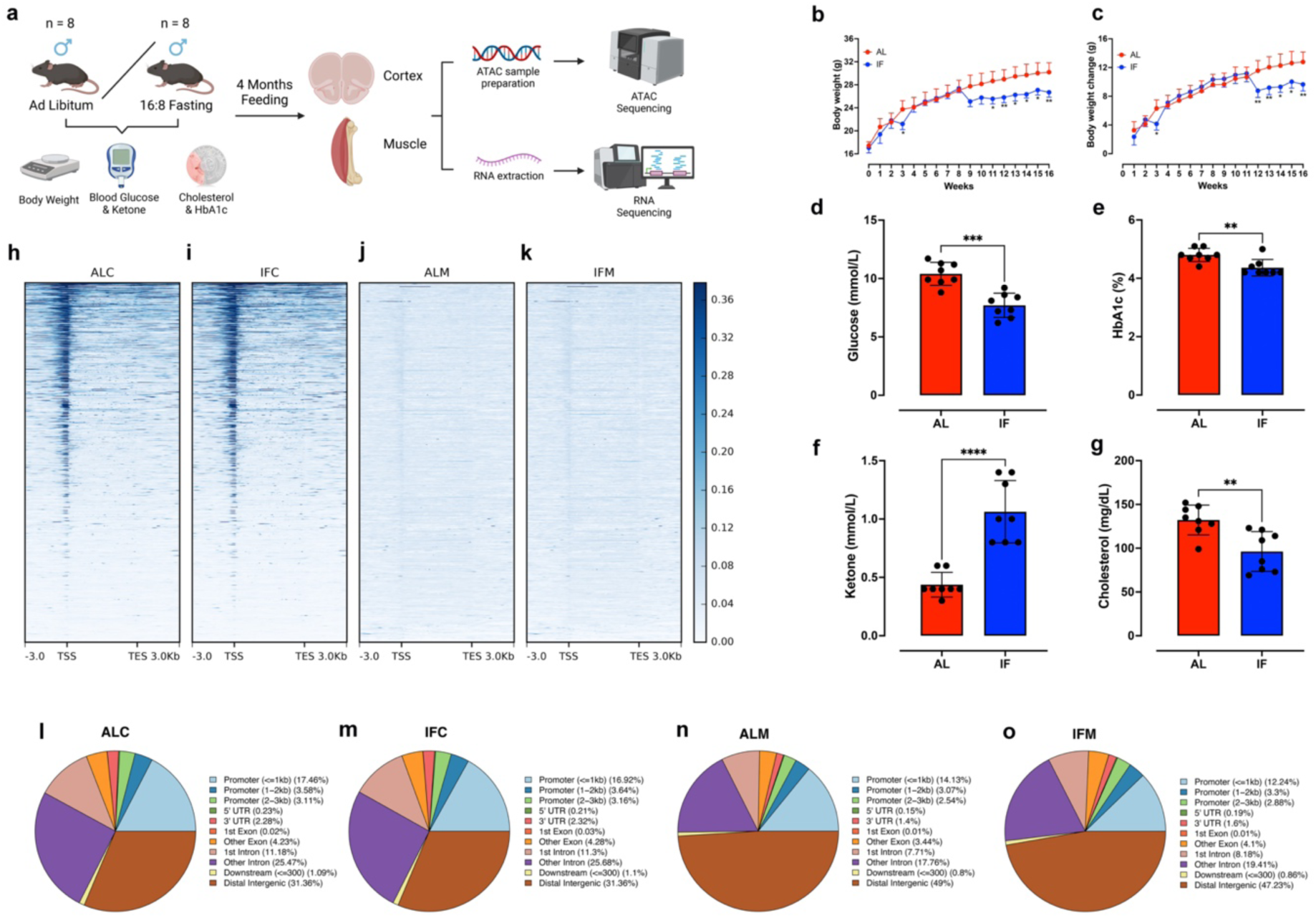
Experimental Design and ATAC-seq. (**a**) Male mice were divided into two groups: Ad Libitum (AL) and 16:8 fasting (IF) (n=8 per group). After 4 months on their respective dietary regimens, mice were sacrificed, and tissues from the cortex and muscle were harvested, prepared, and subjected to ATAC-seq and RNA-seq analyses. (**b-c**) Average weekly body weight and (**b**) body weight changes (**c**) of AL and IF mice over the 16-week experimental period. Statistical analysis was performed using two-way ANOVA with Tukey’s post hoc test. Values represent the mean ± SEM (n=8 mice per group). Significance levels: *p<0.05, **p<0.01 compared to the AL group. Metabolic parameters measured at week 16: fasting blood glucose (**d**), HbA1c levels (**e**), ketone levels (**f**), and cholesterol levels (**g**). Values represent the mean ± SEM (n=8 mice per group). Statistical significance was determined using two-tailed paired T-test with the following significance levels: **p<0.01, ***p<0.001, ****p<0.0001 compared to the AL group. (**h-k**) Heatmaps depicting read distribution mapped to the gene body in AL and IF cortex (**h, i**) and AL and IF muscle groups (**j, k**). The color scale represents fold enrichment values. (**l-o**) Pie charts illustrating the distribution of peaks in functional gene regions for AL and IF groups.

### Evaluation of ATAC-Seq Data Quality

The quality control analysis of ATAC-seq data was conducted to ensure the reliability and accuracy of the sequencing results (**Supplementary Figures 1-2**). The typical fragment size distribution plots for AL and IF cortex tissues (**Supplementary Figure 1a-b**) and muscle tissues (**Supplementary Figure 1c-d**) show enrichment around 100 bp and 200 bp, indicating the presence of nucleosome-free and mono-nucleosome-bound fragments. Insets in **Supplementary Figure 1a-d** reveal that nucleosome-free fragments are enriched at transcription start sites (TSS). Correlation analyses demonstrate a high correlation between biological replicates for both cortex (**Supplementary Figure 1e**) and muscle samples (**Supplementary Figure 1f**).

The successful detection of accessible regions is further supported by the strong enrichment of ATAC-seq reads around TSS sites in cortex samples (**Figure 1h-i**) and moderate enrichment in muscle samples (**Figure 1j-k**). The peak annotation pie chart illustrates that over half of the peaks correspond to enhancer regions (distal intergenic and intronic regions), with approximately 20-25% located in promoter regions for cortex samples in both AL (**Figure 1l**) and IF (**Figure 1m**) conditions. In contrast, nearly 70% of the peaks in muscle samples fall within enhancer regions, with fewer than 20% in promoter regions (**Figure 1n-o**). Overall, the quality control metrics confirm that the ATAC-seq data for AL and IF cortex is robust and suitable for downstream analysis. The high peak density observed across samples, particularly at TSS regions, indicates successful enrichment of accessible chromatin regions, which is critical for accurately interpreting the chromatin landscape. Although muscle samples exhibited lower peak density, the data were sufficient to detect IF-induced changes in chromosome accessibility.

### Mapping of the Open Chromatin Accessibility in Response to Intermittent Fasting

Genome-wide chromatin accessibility in cortex and skeletal muscle tissue samples obtained from AL and IF animals. Cortex and muscle samples in each group (n=8) were independently subjected to ATAC-seq. The ATAC-seq libraries were sequenced, yielding an average of 697 million reads in cortex (43.1 million per sample) and total 676 million reads in muscle (43.61 million per sample). To assess whether IF leads to differential accessibility peaks compared to AL, we compared the log2 fold change in chromatin accessibility peaks between the IF and AL groups (**Figure 2**). The data indicate that IF led to increased chromatin accessibility at certain sites, while decreasing accessibility at other sites within the cortex (**Figure 2a**) and muscle (**Figure 2b**) compared to AL. We utilized Reads Per Million (RPM) values for peaks in different samples (representing the ratio of 1 million reads that enrich a peak in a single sample) to conduct clustering analysis, determining the enrichment pattern of the same peak across cortex (**Figure 2c**) and muscle (**Figure 2d**) samples. Hierarchical clustering based on fold enrichment in peak values for IF and AL samples demonstrated that IF leads to a distinct enrichment pattern in both organs, with different colors representing various clustering groups. We identified differential binding sites by finding peaks with a FoldEnrich ratio greater than 2. Genes associated with these differential sites were annotated, and enrichment analysis was conducted. The Venn diagrams comparing differential peaks in the cortex (**Figure 2e**) and muscle (**Figure 2f**) illustrate the overlap between the AL and IF groups, with red indicating IF and blue indicating AL. The numbers in the pie charts represent peak counts for each group, while the overlapping sections denote common peaks between the comparison groups. Next, we conducted KEGG pathway enrichment analysis for genes related to differential peaks between IF and AL in both cortex (**Figure 2g**) and muscle (**Figure 2h**). The enrichment analysis revealed that IF led to the enrichment of several common pathways, including thermogenesis, ribosome biogenesis, Rap1 signaling, metabolic pathways, HIF-1 signaling, glycolysis, and amino acid biosynthesis (**Figure 2g-h**). To further explore these pathway enrichments, we analyzed the genes involved in selected pathways such as ribosome biogenesis, metabolism, HIF-1 signaling, glycolysis, and amino acid biosynthesis in both cortex (**Figure 2i**) and muscle (**Figure 2j**). These pathways were selected based on their enrichment scores, which exceeded 0.1 in both the cortex and muscle. The metabolic pathway, although not meeting this criterion, was included due to its significant gene count (over 30 in the cortex and over 50 in the muscle) and its strong correlation with dietary intake. In the cortex of IF mice, we observed an enrichment of peaks close to the TSS of genes involved in metabolic pathways such as *Phospholipase A2 Group III (Pla2g3)*, *Prostaglandin D2 Synthase (Ptgds)*, *ATPase H+ Transporting V0 Subunit E (Atp6v0e)*, *Phosphatidylethanolamine N-Methyltransferase (Pemt)*, *TRIT1 tRNA Dimethylallyltransferase (Trit)*, and *Acetyl-CoA Carboxylase Beta (Acacb)*; enrichment of peaks close to the TSS of genes in HIF-1 pathways such as *Phosphoinositide-3-Kinase Regulatory Subunit 1 (Pik3r1)*, *Protein Kinase C Beta (Prkcb)*, *Insulin-Like Growth Factor 1 (Igf1)*; and enrichment of peaks close to the TSS of genes in ribosome-related pathways such as *Signal Recognition Particle 9 (Srp9)* (**Figure 2i**). Additionally, more peaks were observed close to the TSS of noncoding genes, particularly within the ribosome KEGG pathway (**Supplementary Figure 3**). Similarly, in the muscle of IF mice, we observed an enrichment of peaks close to the TSS of genes involved in metabolism, such as *Phospholipase A2 Group VI (Pla2g6)*, *Cytochrome C Oxidase Subunit 10 (Cox10)*, *Nudix Hydrolase 2 (Nudt2)*, *Selenophosphate Synthetase (Sephs)*, and *Insulin 1 (Ins1)*; and an enrichment of peaks close to the TSS of genes in HIF-1 pathways such as *Muscle Phosphofructokinase (Pfkm)* (**Figure 2j**). The heat map for all peaks near the TSS of genes within the ribosome, metabolism, HIF-1, and glycolysis KEGG pathways is presented in **Supplementary Figure 4**. Furthermore, we conducted an analysis of Gene Ontology (GO) term pathways for the enrichment of peaks close to the TSS of genes in both the cortex and muscle (**Supplementary Figure 5**). This analysis included GO terms associated with downregulated peaks (**Supplementary Figure 5a and c**), and GO terms associated with upregulated peaks (**Supplementary Figure 5b and d**) in the cortex and muscle in response to IF.

**Figure 2:**
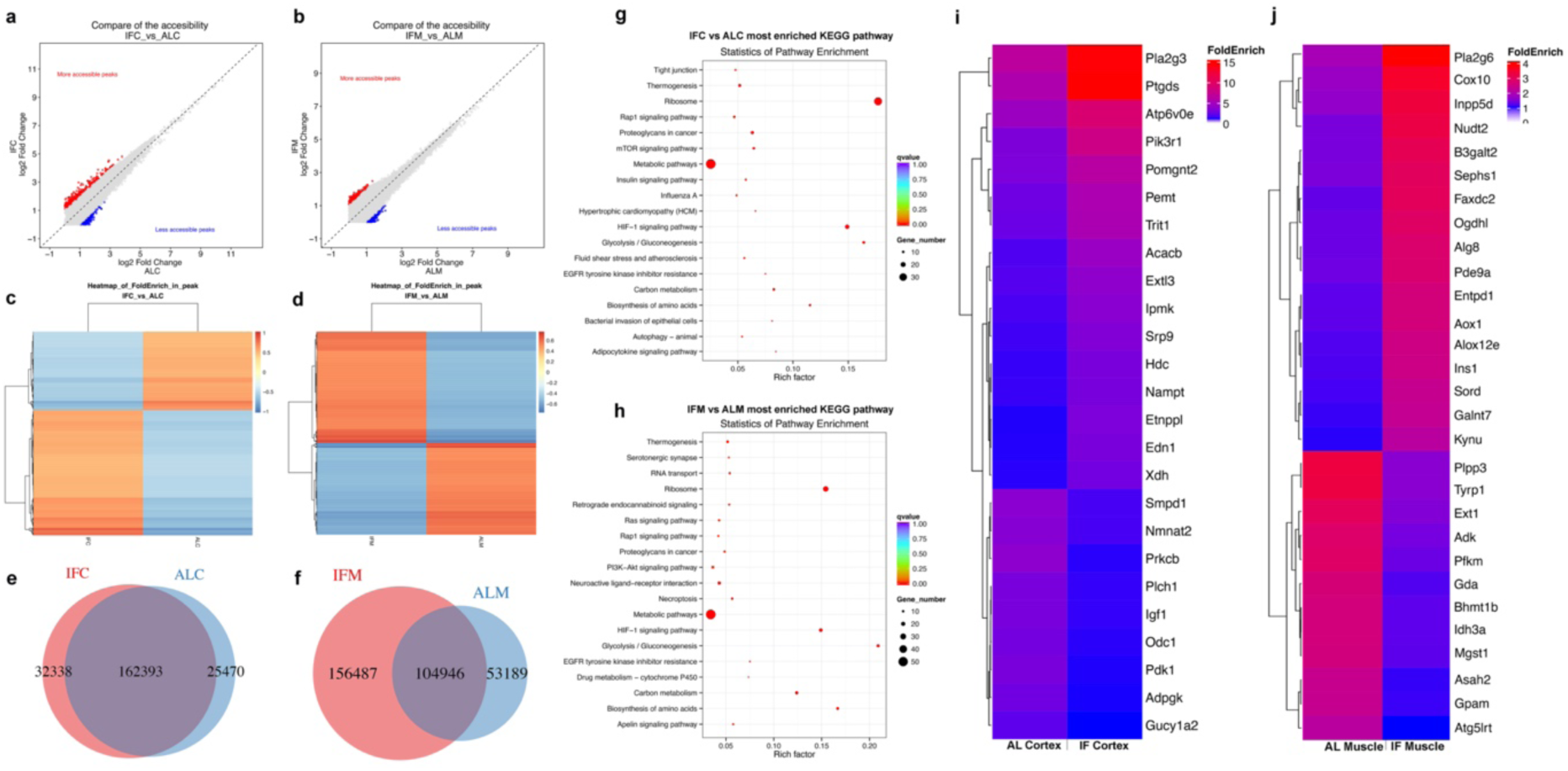
Mapping Open Chromatin Accessibility in Response to Intermittent Fasting. (**a and b**) Comparison of chromatin accessibility peaks between IF (intermittent fasting) and AL (ad libitum) in the cortex (**a**) and muscle (**b**) tissues using log2 fold change. Each dot represents a peak, with a log2 fold change ≥1 considered significant. (**c and d**) Heatmaps illustrating the fold enrichment of peaks in the cortex (**c**) and muscle (**d**) tissues between IF and AL. Each horizontal line represents a peak, with color intensity indicating the degree of fold enrichment. (**e and f**) Venn diagrams showing the number of common and differential peaks between IF and AL in the cortex (**e**) and muscle (**f**). (**g and h**) Scatter plots depicting the most enriched KEGG pathways in the cortex (**g**) and muscle (**h**) in response to IF versus AL. The size of each dot represents the number of overlapping genes within the pathway. (**i and j**) Heatmaps displaying the fold enrichment of genes associated with significantly enriched pathways in the cortex (**i**) and muscle (**j**). These pathways include ribosome biogenesis, metabolism, HIF-1 signaling, and glycolysis. The color scale represents fold enrichment values, with red indicating higher and blue indicating lower fold enrichment. Pathways were selected based on q-values (q-value < 0.05).

### Analysis of Transcription Factor Binding Motifs in Response to Intermittent Fasting

The motif indicates the sequence conservation at the peak position, which may play a critical role in the regulation of gene expression. Using sequence information from 250bp upstream and downstream of the peak (totaling 500bp), we employed Homer software to identify conserved sequence features of peak enrichment (**Figure 3**). To identify DNA-binding transcription factors (TFs) that link differential chromatin accessibility and gene expression, motif analysis was performed. We identified the top 10 highly-expressed known TFs with enriched motifs in distal regions exhibiting increased accessibility in response to IF in the cortex (**Figure 3a-b**) and muscle (**Figure 3c-d**) compared to AL conditions. The TFs identified in the cortex are: Sp1 (Specificity Protein 1), Nfia (Nuclear Factor I A), Nfil3 (Nuclear Factor Interleukin 3 Regulator), Jun (Jun Proto-Oncogene), Mef2a (Myocyte Enhancer Factor 2A), Zfp189 (Zinc Finger Protein 189), Neurod2 (Neurogenic Differentiation 2), Elf1 (E74 Like ETS Transcription Factor 1), Pbx2 (Pre-B Cell Leukemia Homeobox 2), and Banp (B-Cell Activation Factor Nuclear Factor) (**Figure 3a**). The Integrative Genomics Viewer (IGV) demonstrates peaks corresponding to these TFs in response to IF compared to AL (**Figure 3b**). The top 10 TFs enriched in muscle include Samd4 (Sterile Alpha Motif Domain Containing 4), Zeb2 (Zinc Finger E-Box Binding Homeobox 2), Tcf4 (Transcription Factor 4), Stat5b (Signal Transducer and Activator of Transcription 5B), Rest (RE1 Silencing Transcription Factor), Nkx3-1 (NK3 Homeobox 1), Ptbp1 (Polypyrimidine Tract Binding Protein 1), Zbtb7b (Zinc Finger and BTB Domain Containing 7B), Gata5 (GATA Binding Protein 5), and Snrnp70 (Small Nuclear Ribonucleoprotein U70) (**Figure 3c**). IGV in Figure 3D shows peaks corresponding to these TFs in response to IF compared to AL in muscle (**Figure 3d**).

**Figure 3:**
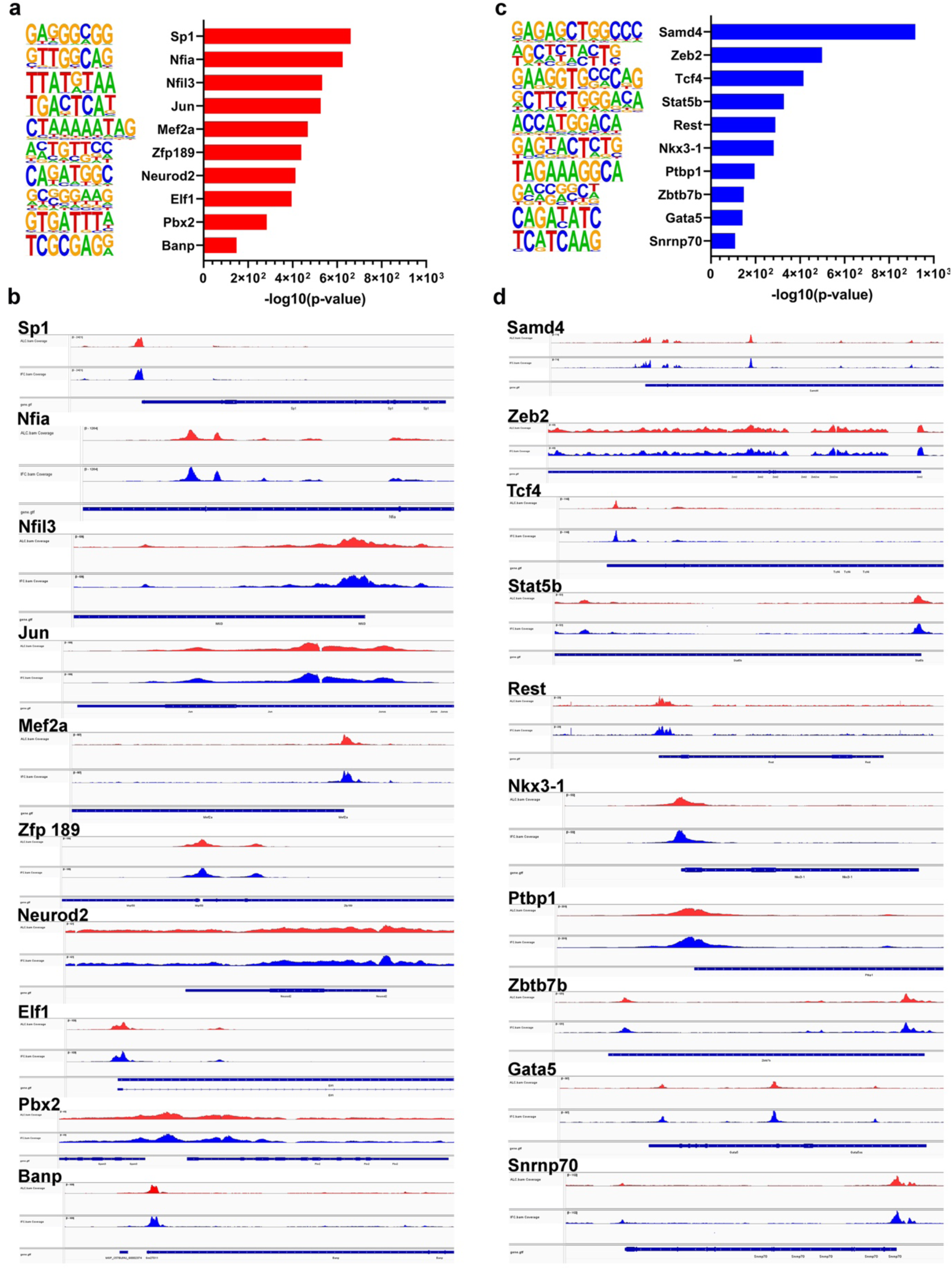
Analysis of Transcription Factor Binding Motifs in Response to Intermittent Fasting. Top 10 most enriched transcription factors (TFs) and their corresponding motif logos in IF compared to AL in the cortex (**a and b**) and muscle (**c and d**). Transcription factors are ranked according to their -log10 p-value. Visualization of the top 10 most enriched transcription factors in IF compared to AL in the cortex (**b**) and muscle (**d**) using Integrative Genomics Viewer. Red tracks represent AL, and blue tracks represent IF. The track labeled “gene.gtf” shows the gene names, with arrows on the gene body indicating the direction of transcription. The height of the peaks represents the number of reads.

While we have compared TFs with enriched motifs in both cortex and muscle between IF and AL conditions, we also analyzed TFs with enriched motifs in IF samples against baseline data (**Supplementary Figure 6**). Subsequently, we performed IGV analysis of these identified TFs in IF tissues compared to AL tissues. Some of these TFs were also observed in our direct IF vs AL comparison. However, new TFs have emerged, potentially providing further insights into the transcriptional mechanisms underlying the effects of IF. These newly enriched TFs in the cortex include NR3C2 (Nuclear Receptor Subfamily 3 Group C Member 2), SRSF1 (Serine And Arginine Rich Splicing Factor 1), TEAD4 (TEA Domain Transcription Factor 4), EOMES (Eomesodermin), ELK4 (ETS Like-1 Protein 4), YY1 (Yin Yang 1), HNF4A (Hepatocyte Nuclear Factor 4 Alpha), BCL6B (B Cell CLL/Lymphoma 6B), and IRF3 (Interferon Regulatory Factor 3) (**Supplementary Figure 6a**), with corresponding IGV analysis comparing these TFs in IF cortex to AL cortex samples (**Supplementary Figure 6b**). Similarly, TFs with enriched motifs in muscle IF samples against baseline include STAT1 (Signal Transducer and Activator of Transcription 1), HOXC10 (Homeobox C10), POU4F3 (POU Class 4 Homeobox 3), ENOX1 (Ecto-NOX Disulfide-Thiol Exchanger 1), KHDRBS1 (KH RNA Binding Domain Containing Signal Transduction Associated 1), ZFP410 (Zinc Finger Protein 410), SF3B4 (Splicing Factor 3B Subunit 4), MSX3 (Msh Homeobox 3), SMAD2 (SMAD Family Member 2), and LIN28A (Lin-28 Homolog A) (**Supplementary Figure 6c**). IGV analysis illustrating these TFs in IF muscle compared to AL muscle samples is shown in **Supplementary Figure 6d**. The enrichment in both organs suggests a regulatory role for these TFs in chromatin remodeling and gene expression modulation in response to IF. These findings provide new insights into the transcriptional mechanisms underlying the response of brain and muscle cells to dietary interventions.

### Integration of ATAC-seq and RNA-seq Results

To investigate the relationship between alterations in chromatin accessibility and changes in gene expression, we conducted an integrative analysis of RNA-Seq and ATAC-Seq data. RNA-Seq analysis of cortical tissues revealed 4,159 differentially expressed genes in response to IF (**Figure 4a**). GO terms analysis of cortex RNA-seq indicated significant changes in genes associated with cognition, learning and memory, positive regulation of neuronal differentiation, positive regulation of neural migration, and regulation of synapse structure following IF (**Figure 4b**). Additionally, we assessed the cellular components and molecular functions that were significantly affected by IF in the cortex. Our data support the notion that IF enhances neuronal cellular components, such as synapses, synaptic membranes, and integral components of the postsynaptic membrane (**Supplementary Figure 7a-b**). Molecular function analysis of cortex revealed that IF promotes GTPase regulator activity, ion channel activity, ion transmembrane transporter activity, and calmodulin binding (**Supplementary Figure 7c-d**).

**Figure 4:**
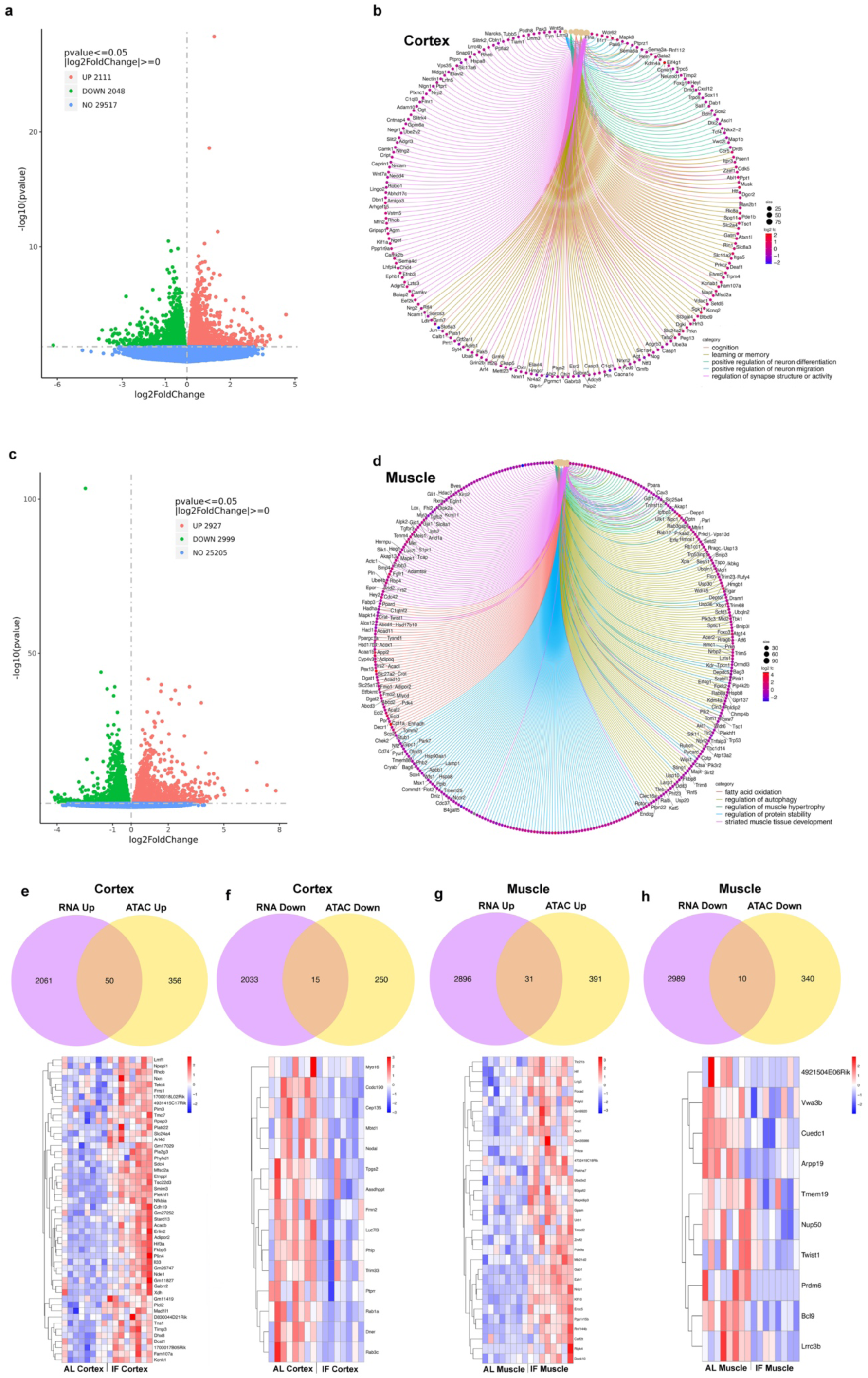
Integrative analysis of ATAC-seq and RNA-seq Data. (**a and c**) Volcano plots of differentially expressed genes for the IF groups compared to the AL group in both cortex (**a**) and muscle (**c**). The threshold for differential expression was set at p<0.05. Each dot represents an individual gene (blue: no significant difference; red: upregulated gene; green: downregulated gene). The volcano plots display statistical significance (-log10 p-value) against enrichment (log2 fold change) of differentially expressed genes. (**b and d**) Circular network plots (cnetplot) showing significantly enriched Gene Ontology (GO) terms from the Biological Process (BP) category for the IF group compared to the AL group in both cortex (**b**) and muscle (**d**). Each node represents a gene involved in the enriched GO terms. Nodes are colored according to the log2 fold change (log2FC) of the corresponding FPKM values, with red indicating upregulation and blue indicating downregulation. Terms were selected based on adjusted p-values (adjusted p < 0.05). (**e-h**) Venn diagrams illustrating the common and differentially expressed genes between RNA-seq (upregulated/downregulated) and ATAC-seq (upregulated/downregulated) in the cortex (**e and f**) and muscle (**g and h**). Heatmaps of commonly expressed genes in RNA-seq data from the cortex and muscle are displayed below the Venn diagrams. The colour scale represents z-score values, with red indicating upregulation and blue indicating downregulation.

RNA-Seq analysis of muscle tissues identified 5,926 differentially expressed genes in response to IF (**Figure 4c**). Cnetplot analysis showed that genes involved in fatty acid oxidation, regulation of autophagy, regulation of muscle hypertrophy, regulation of protein stability, and muscle tissue development were significantly altered following IF (**Figure 4d**). Furthermore, we analyzed cellular components and molecular functions in muscle tissues. IF was found to enhance the extracellular matrix, mitochondrial function, and both mitochondrial and organelle inner membranes (**Supplementary Figure 8a-b**). Molecular function analysis demonstrated that IF significantly affects insulin receptor substrate binding, nuclear hormone receptor binding, transcription factor binding, and transcription co-regulator activity, among other functions (**Supplementary Figure 8c-d**).

Next, we integrated the differentially expressed genes from RNA-Seq with the genes exhibiting altered chromatin accessibility in both organs (**Figure 4e–h and Supplementary Figure 9**). Our analysis identified 50 common upregulated genes and 15 common downregulated genes in the cortex in response to IF (**Figure 4e-f**). Similarly, in muscle tissue, we found 31 common upregulated genes and 10 common downregulated genes associated with IF (**Figure 4g-h**). To visualize these findings, we created heatmaps for the commonly upregulated and downregulated genes in both organs (**Figure 4e-h**). Commonly upregulated genes in the cortex in response to IF include *Lipid Metabolism Factor 1 (Lmf1)*, *Ras Homolog Family Member B (Rhob)*, *Tektin 4 (Tekt4)*, *Ferroxidase 1 (Frrs1)*, *Proto-Oncogene Serine/Threonine-Protein Kinase Pim-3 (Pim3)*, *Transmembrane Channel-Like 7 (Tmc7)*, *Syndecan 4 (Sdc4)*, *Nuclear Factor Kappa B Inhibitor Alpha (Nfkbia)*, *Acetyl-CoA Carboxylase Beta (Acacb)*, *Tissue Inhibitor of Metalloproteinases 3 (Timp3)*, and *Potassium Channel Subfamily K Member 1 (Kcnk1)*, among others (**Figure 4e**). On the other hand, genes downregulated in response to IF include *Myosin XVI (My016)*, *Coiled-Coil Domain Containing 190 (Ccdc190)*, *Centrosomal Protein 135 (Cep135)*, *Nodal Growth Differentiation Factor (Nodal)*, *Mbt Domain-Containing 1 (Mbtd1)*, *PH-Interacting Protein (Phip)*, and *Ras-Related Protein Rab-1A (Rab1a)*, among others (**Figure 4f**). Genes that are commonly upregulated in response to IF in muscle include *Tetratricopeptide Repeat Domain 21B (Ttc21b)*, *Hypoxia-Inducible Factor (Hif)*, *Aldehyde Oxidase 1 (Aox1)*, *Fibroblast Growth Factor Receptor Substrate 2 (Frs2)*, *Glycerol-3-Phosphate Acyltransferase, Mitochondrial (Gpam)*, *Zinc and Ring Finger 2 (Znrf2)*, *Phosphodiesterase 9A (Pde9a)*, *Excision Repair Cross-Complementation Group 5 (Ercc5)*, *Kruppel-Like Factor 10 (Klf10)*, *Nuclear Receptor Interacting Protein (Nrip)*, *Enhancer of Zeste Homolog 1 (Ezh1)*, and *GRB2-Associated Binding Protein 1 (Gab1)*, among others (**Figure 4g**). Conversely, genes commonly downregulated in response to IF in muscle include *PR/SET Domain 6 (Prdm6)*, *B-Cell CLL/Lymphoma 9 (Bcl9)*, *Leucine Rich Repeat Containing 3B (Lrrc3b)*, *Twist Family BHLH Transcription Factor 1 (Twist1)*, and *cAMP-Regulated Phosphoprotein 19 (Arpp19)*, among others (**Figure 4h**).

## Discussion

IF is acclaimed for its numerous health benefits in both animals and humans, protecting against neurological diseases, cardiovascular conditions, and metabolic disorders, while also extending healthspan and lifespan (25). This study significantly advances our understanding the mechanisms of IF’s health benefits by demonstrating for the first time that IF can alter chromosome accessibility and profoundly influence gene expression and phenotypic outcomes.

Previously, research from our group has revealed that a 16-hour IF regimen provides protection against stroke and vascular dementia, notably influencing gene expression across multiple pathways in the cortex, cerebellum, heart, and liver (4, 9, 12). Gene expression is regulated by a complex network of mechanisms. To understand how IF regulates gene expression, we have examined epigenetic mechanisms and found that IF modifies the DNA methylation landscape and histone modification to influence IF-induced gene expression (4). While other epigenomic modifications may be modulated in response to IF, studies have established how IF affects H3K9 trimethylation and DNA methylation to regulate myriad metabolic processes, bring about robust metabolic switching responses in the brain, and protect against dementia (19). Furthermore, proteomics and phospho-proteomics analyses illuminate the complex molecular mechanisms such as cyclic GMP signaling through which IF confers health benefits and aids disease prevention (20). A comparative analysis of the correlation between transcript and protein abundance following IF revealed a correlation, suggesting that IF-induced gene expression and its translation are pivotal in delivering the phenotypic benefits observed (20). Collectively, these previous studies provide valuable insights into how IF-induced metabolic switching can alter epigenetic, genetic, and proteomic mechanisms to modify biological pathways and remodel tissues and organs to benefit organisms. However, whether metabolic switching by IF leads to changes in chromosome accessibility to influence gene expression remains unanswered. Here, we show that IF leads to differences in chromosomal availability, which potentially contributes to gene expression regulation in brain and muscle tissues.

Analyzing the differential peaks in response to IF versus AL feeding revealed that genes associated with these peaks are modifying critical biological pathways. These pathways include ribosome biogenesis, metabolism, thermogenesis, HIF signaling, glycolysis, carbon metabolism, and the synthesis of amino acids in both the cortex and muscle. Additionally, autophagy pathway was modulated in the cortex, while PI3K-Akt, Ras, and Rap1 signaling pathways were influenced in muscle tissue. Our deeper investigation into these genes associated with these peaks demonstrated that IF-induced changes in chromosome accessibility form opportunities for transcription factors to initiate the expression of genes that could be pivotal in mediating the health benefits associated with IF. We identified several transcription factors with significantly higher motifs in the IF cortex compared to the AL cortex. These transcription factors include Specificity Protein 1 (Sp1), which is crucial for cell proliferation, apoptosis, and differentiation (26); Nuclear Factor I/A (NFIA), known for its role in neural development and glial differentiation (27); Nuclear Factor, Interleukin 3 Regulated (Nfil3), involved in circadian rhythm regulation and immune function (28); Jun Proto-Oncogene (Jun), a component of the AP-1 transcription factor complex that regulates cell proliferation and apoptosis (29); Myocyte Enhancer Factor 2A (Mef2a), essential for development and differentiation (30); Zinc Finger Protein 189 (Zfp189), although its specific functions are less well-characterized, it plays a role in gene regulation (31); NeuroD2, important for neuronal differentiation and development (32); E74-Like Factor 1 (Elf1), which regulates gene expression, particularly in immune cells (33); and Brain and Nervous System-Associated Proto-Oncogene (Banp), associated with neural development and potentially playing a role in neurogenesis (34). These transcription factors are pivotal in understanding how IF-induced changes in chromosome accessibility can lead to the initiation of gene expression in the brain.

Similarly, analysis in skeletal muscle revealed several transcription factors with significantly higher motifs in the IF group compared to the AL group. These include SMAD Family Member 4 (SMAD4), which is involved in muscle differentiation and regeneration through the TGF-beta signaling pathway (35). Zinc Finger E-Box Binding Homeobox 2 (ZEB2) is known to play a role in muscle satellite cell quiescence and differentiation (36). Transcription Factor 4 (TCF4) is implicated in the maintenance of muscle stem cells and muscle regeneration (37). Signal Transducer and Activator of Transcription 5B (STAT5B) contributes to muscle growth and repair through cytokine signaling (38). RE1-Silencing Transcription Factor (REST) has been shown to influence muscle fiber type specification and neuromuscular junction maintenance (39). NK3 Homeobox 1 (NKX3-1) is involved in the regulation of muscle cell proliferation and differentiation (40). Polypyrimidine Tract Binding Protein 1 (PTBP1) regulates alternative splicing of pre-mRNAs critical for muscle function and adaptation (41). Zinc Finger and BTB Domain Containing 7B (ZBTB7B) plays a role in muscle development and also forms a ribonucleoprotein transcriptional complex with the lncRNA Blnc1 and drives thermogenic gene expression (42). GATA Binding Protein 5 (GATA5) is involved in muscle development and function, influencing muscle cell differentiation (43). Lastly, Small Nuclear Ribonucleoprotein U1 Subunit 70 (SNRNP70) is essential for the splicing of pre-mRNAs, which is critical for gene expression and the regulation of synapse formation and neuromuscular synaptogenesis (44). These transcription factors highlight the intricate molecular mechanisms through which IF influences gene expression and enhances muscle health and function.

While our data show that IF led to changes in chromosome accessibility, it is crucial to understand whether these changes also result in gene expression in both the cortex and muscle. To address this, we performed RNA sequencing on the same tissues and conducted integrative analyses. Consistent with our previous findings, RNA sequencing revealed that IF induced the expression of genes involved in pathways such as calcium signaling, PI3K-Akt signaling, circadian rhythm, metabolism, axon guidance, MAPK signaling, and synaptic pathways in the cortex. In the muscle, analysis of differentially expressed genes showed that IF activated pathways related to thermogenesis, focal adhesion, insulin signaling, neurotrophin signaling, and pluripotency, among others. Furthermore, we used cnetplot functions to gain a deeper understanding of gene expression changes in the cortex and muscle in response to IF compared to AL. Cnetplot analysis identified numerous genes that may contribute to the health benefits observed with IF.

Integrative analysis of differentially expressed genes from RNA sequencing and enrichment of peaks of genes close to the TSS in ATAC-seq revealed 50 upregulated and 15 downregulated common genes in response to IF in the cortex and 31 upregulated and 10 downregulated genes in response to IF in muscle compared to AL. In the cortex, highly significant upregulated genes include KCNK1 (Potassium Channel, Subfamily K, Member 1), also known as TWIK-1 involved in maintaining the resting membrane potential (45); TIMP3 (Tissue Inhibitor of Metalloproteinases 3), which plays a role in inhibiting metalloproteinases and influencing tissue remodeling (46); PLEKHF1 (Pleckstrin Homology And FYVE Domain Containing 1), involved in intracellular signaling (47); SDC4 (Syndecan 4), which plays a role in formation of focal adhesions and interacts with many growth factors to regulate cell migration and neural induction (48); and TEKT4 (Tektin 4), associated with cytoskeletal structure (49). Downregulated common genes in the cortex include MYO16 (Myosin XVI), involved in cellular movement and transport (50); MBTD1 (Mbt Domain Containing 1), involved in chromatin regulation (51); RAB1A and RAB3C (Members of RAS Oncogene Family), which plays a role in vesicle transport and a key player in the pathogenesis of Parkinson’s Disease and neurodegeneration (52, 53); DNER (Delta/Notch-Like EGF Repeat Containing), implicated in neural development and signaling (54).

In the muscle, highly upregulated common genes include but are not limited to CSTF2T (Cleavage Stimulation Factor, 3’ Pre-RNA, Subunit 2, Tau Variant), regulates expression of histones and histone-like proteins (55); ERCC5 (Excision Repair Cross-Complementation Group 5), essential for DNA repair (56); EZH1 (Enhancer of Zeste Homolog 1), a part of the Polycomb group proteins that regulate gene expression through chromatin modification (57); PDE9A (Phosphodiesterase 9A), which hydrolyzes cyclic nucleotides and is involved in cellular signaling (58); ZNRF2 (Zinc and Ring Finger 2), playing a role in ubiquitination processes (59); and TTC21B (Tetratricopeptide Repeat Domain 21B), which is crucial for ciliary function and structure (60). Commonly downregulated genes in muscle include BCL9 (B-Cell CLL/Lymphoma 9), involved in Wnt signaling and cell adhesion (61); PRDM6 (PR/SET Domain 6), which functions as a histone methyltransferase (62); TWIST1 (Twist Family BHLH Transcription Factor 1), has been associated with the embryonic development, cancer, and fibrotic diseases, but also the regulation of lipid and glucose metabolism (63); NUP50 (Nucleoporin 50kDa), which is a component of the nuclear pore complex (64); and ARPP19 (CAMP-Regulated Phosphoprotein 19), which has oncogenic functions (65). While further validation is needed to fully comprehend the role of these gene expression changes in response to IF in both the cortex and muscle, our findings validate the changes in chromosome accessibility observed in response to IF. This integrative approach underscores the multifaceted impact of IF on gene expression, highlighting the complex biological pathways through which IF may exert its beneficial effects on health.

In conclusion, this study provides the first evidence that IF alters chromatin accessibility and profoundly influences gene expression. Our findings reveal that IF modifies critical biological and signaling pathways in both the cortex and muscle. By analyzing transcription factors and their motifs, we identified key regulatory elements driving these gene expression changes. Integrative analysis of RNA-Seq and ATAC-Seq data further validated the impact of IF, with numerous genes being upregulated or downregulated in response to IF, highlighting the intricate molecular mechanisms involved. These findings suggest that IF-induced metabolic switching and subsequent changes in chromatin accessibility are central to the health benefits associated with IF. This study underscores the multifaceted impact of IF on gene regulation and elucidates the complex biological pathways through which IF may confer its health benefits.

## Data availability

High throughput ATAC- and RNA-seq data from this manuscript will be submitted to the NCBI Sequence Read Archive (SRA).

## Acknowledgements

Figure 1-Supplementary figure 1 in this article was created using BioRender. This work was supported by La Trobe University (start-up grant to Thiruma V. Arumugam) and the National Health and Medical Research Council of Australia (Grant Identification Number 2019100).

## Animal Ethics

All *in vivo* experimental procedures were approved by La Trobe University (Ethics approval number: AEC21047) Animal Care and Use Committees and performed according to the guidelines set forth by the Australian Code for the Care and Use of Animals for Scientific Purposes (8th edition) and confirmed NIH Guide for the Care and Use of Laboratory Animals.

## Author contributions

Study conception and design: T.V.A., Y.F., X.P., J.G., M.P.M., and D.G.J.; experiment or data collection: Y.F., X.P., N.I.T., X.C., and T.V.A.: data analysis: Y.F., X.P., M.E., and T.V.A.; data interpretation: Y.F., X.P., M.E., Y.K., and T.V.A.; writing-manuscript preparation and intellectual input: Y.F., X.P., Y.K., Y.U.L., G.C., E.O., T.G.J., C.G.S., J.G., M.K.P.L., M.P.M., D.G.J., and T.V.A.; supervision and administration: T.V.A.

## Supplementary Figures and Legends

**Supplementary Figure 1:**
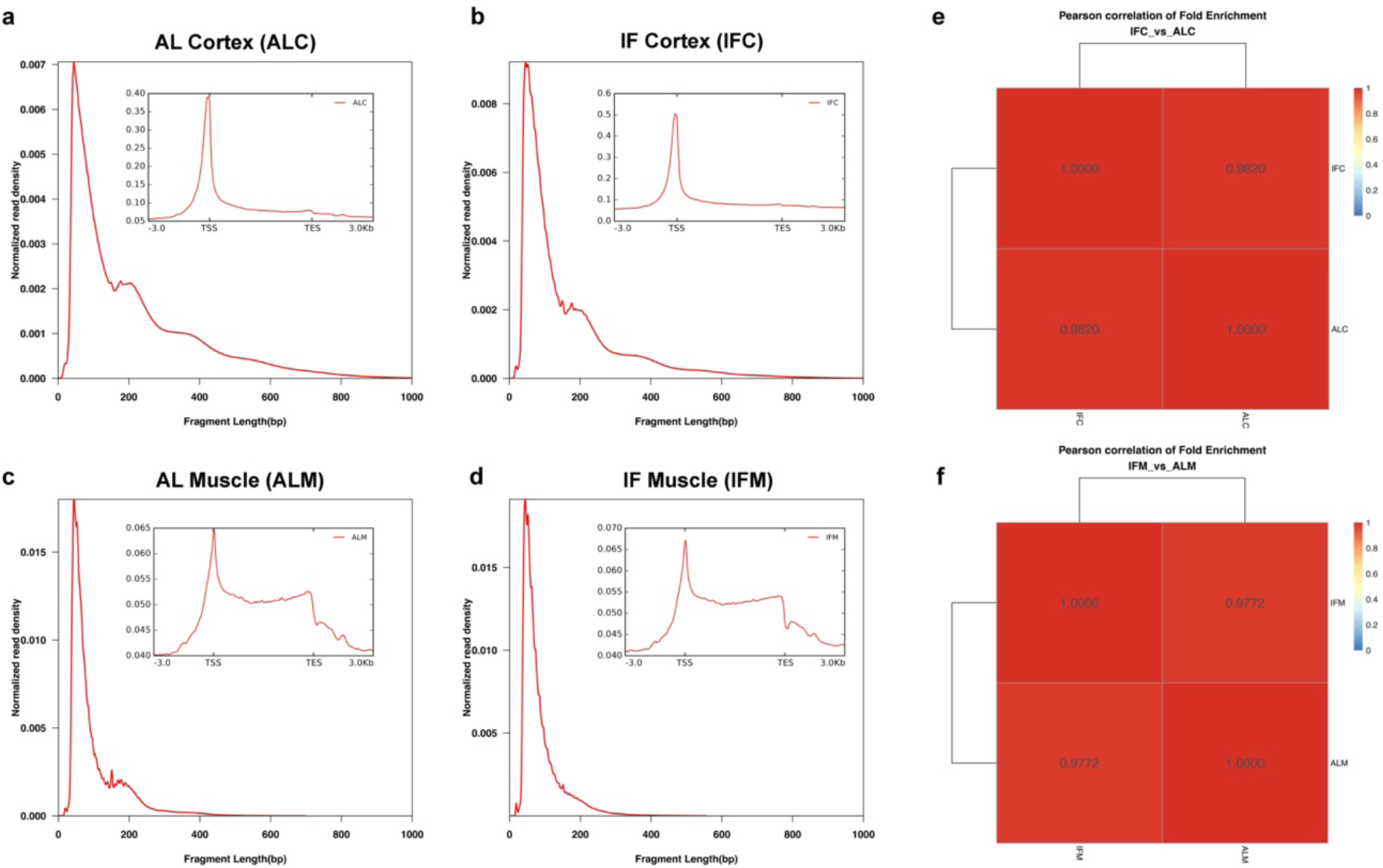
ATAC-Seq Data Quality. (**a and b**) Fragment size distribution plots for cortex tissues from AL (ad libitum) and IF (intermittent fasting) mice. (**c and d**) Fragment size distribution plots for muscle tissues from AL and IF mice. (**e**) Pearson correlation analysis of fold enrichment comparison between AL and IF cortex tissues. (**f**) Pearson correlation analysis of fold enrichment comparison between AL and IF muscle tissues.

**Supplementary Figure 2:**
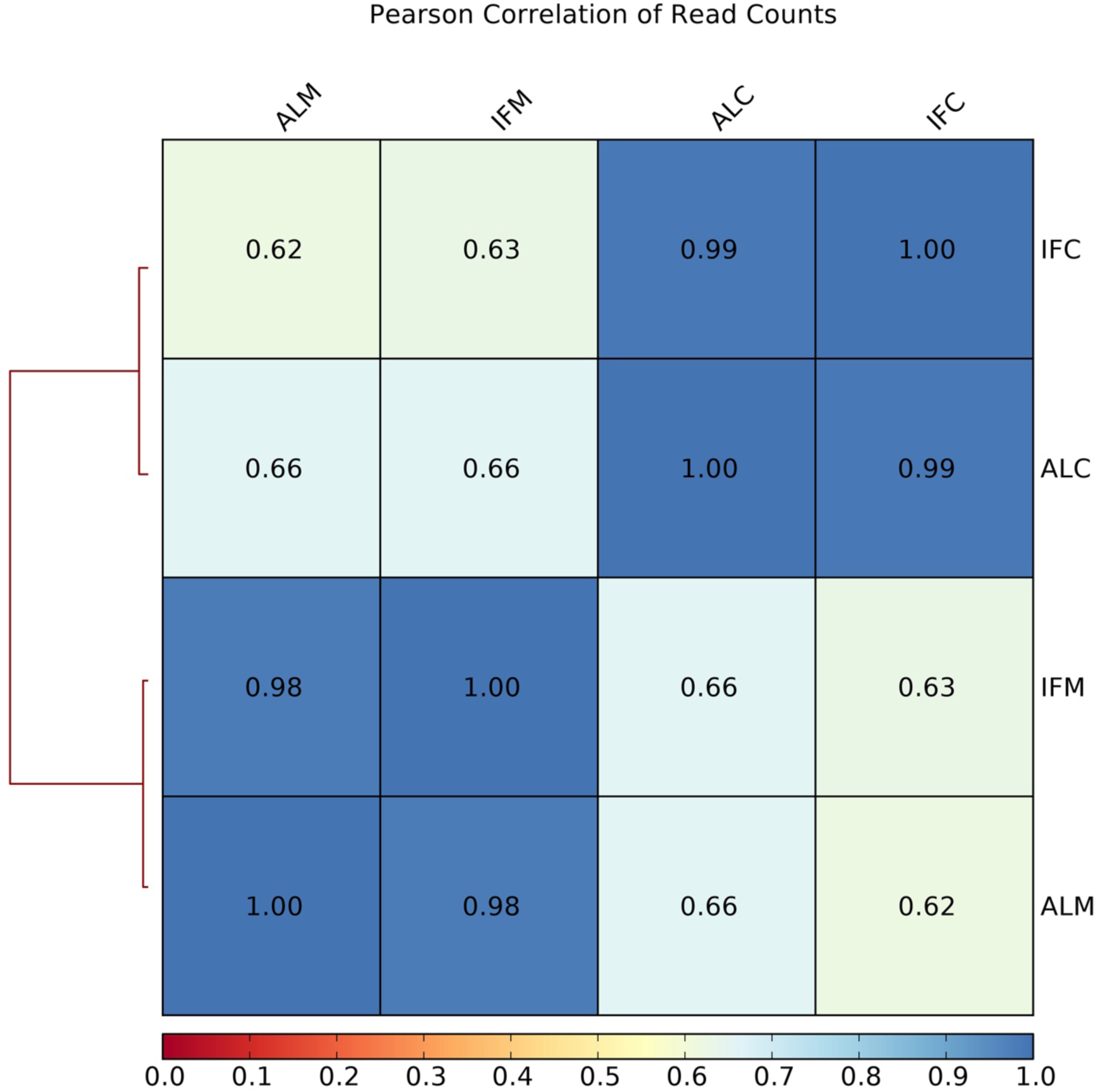
Pearson correlation analysis of fold enrichment comparing AL (ad libitum) and IF (intermittent fasting) in both cortex and muscle tissues.

**Supplementary Figure 3:**
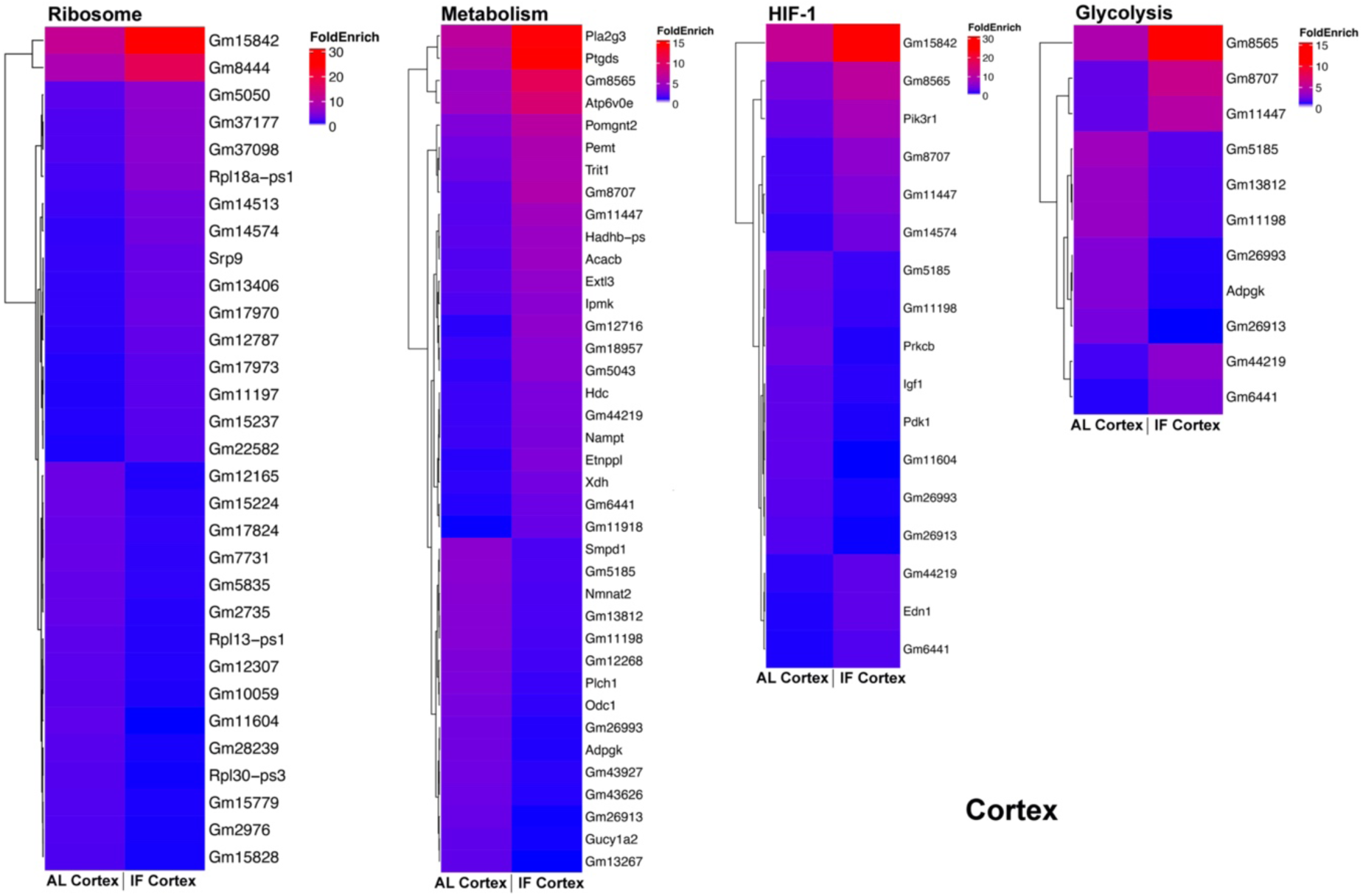
Enriched KEGG Pathway-Related Genes in Cortex from ATAC-seq. Heatmap illustrating the fold enrichment of genes associated with significantly enriched KEGG pathways in the cortex, identified through ATAC-seq. Notable pathways include ribosome biogenesis, metabolism, HIF-1 signaling, and glycolysis. The color scale indicates fold enrichment values, with red representing higher fold enrichment and blue indicating lower fold enrichment. Pathways were selected based on their statistical significance (q-value < 0.05).

**Supplementary Figure 4:**
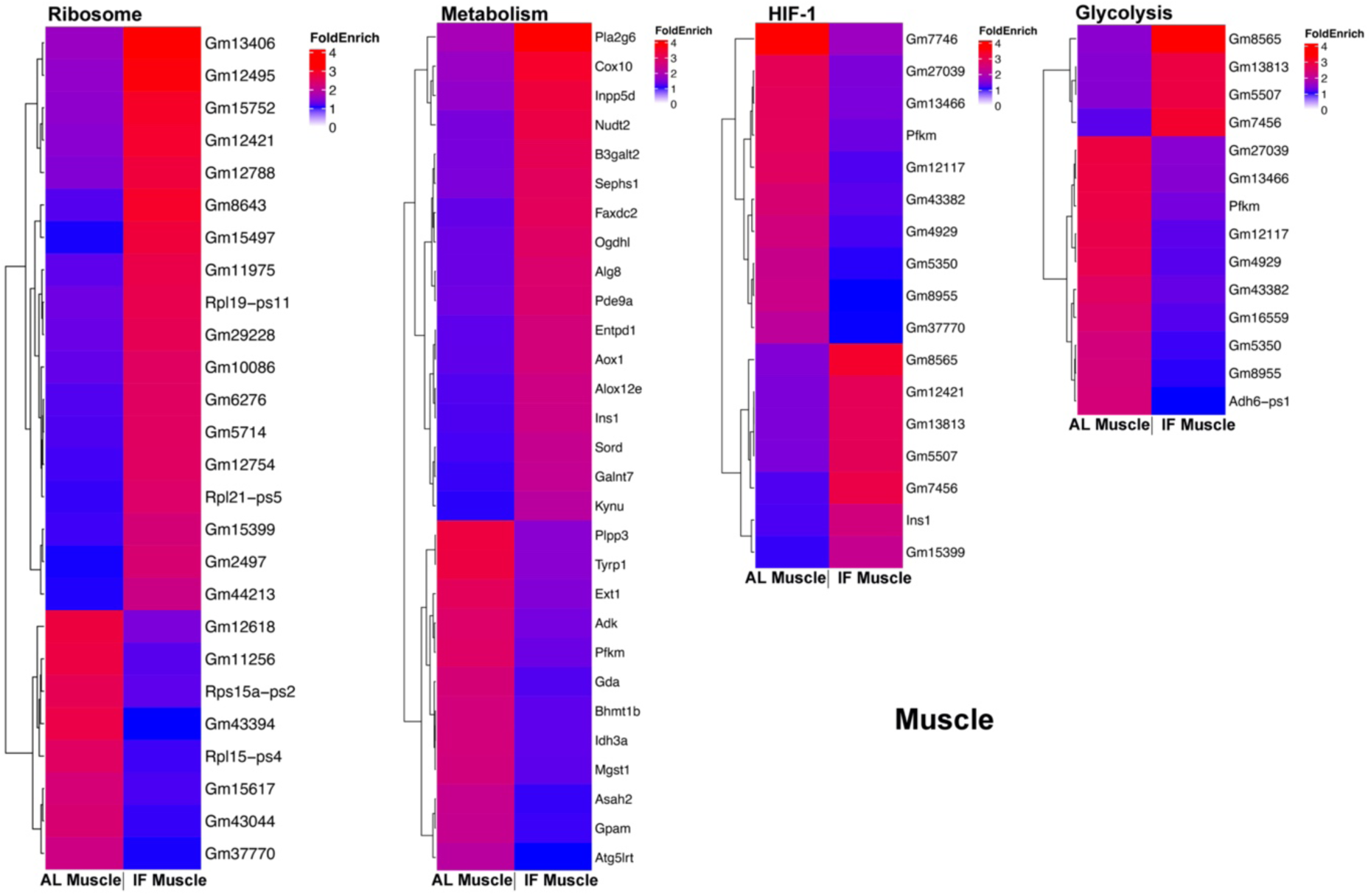
Enriched KEGG Pathway-Related Genes in Muscle from ATAC-seq. Heatmap illustrating the fold enrichment of genes associated with significantly enriched KEGG pathways in the muscle, identified through ATAC-seq. Notable pathways include ribosome biogenesis, metabolism, HIF-1 signaling, and glycolysis. The color scale indicates fold enrichment values, with red representing higher fold enrichment and blue indicating lower fold enrichment. Pathways were selected based on their statistical significance (q-value < 0.05).

**Supplementary Figure 5:**
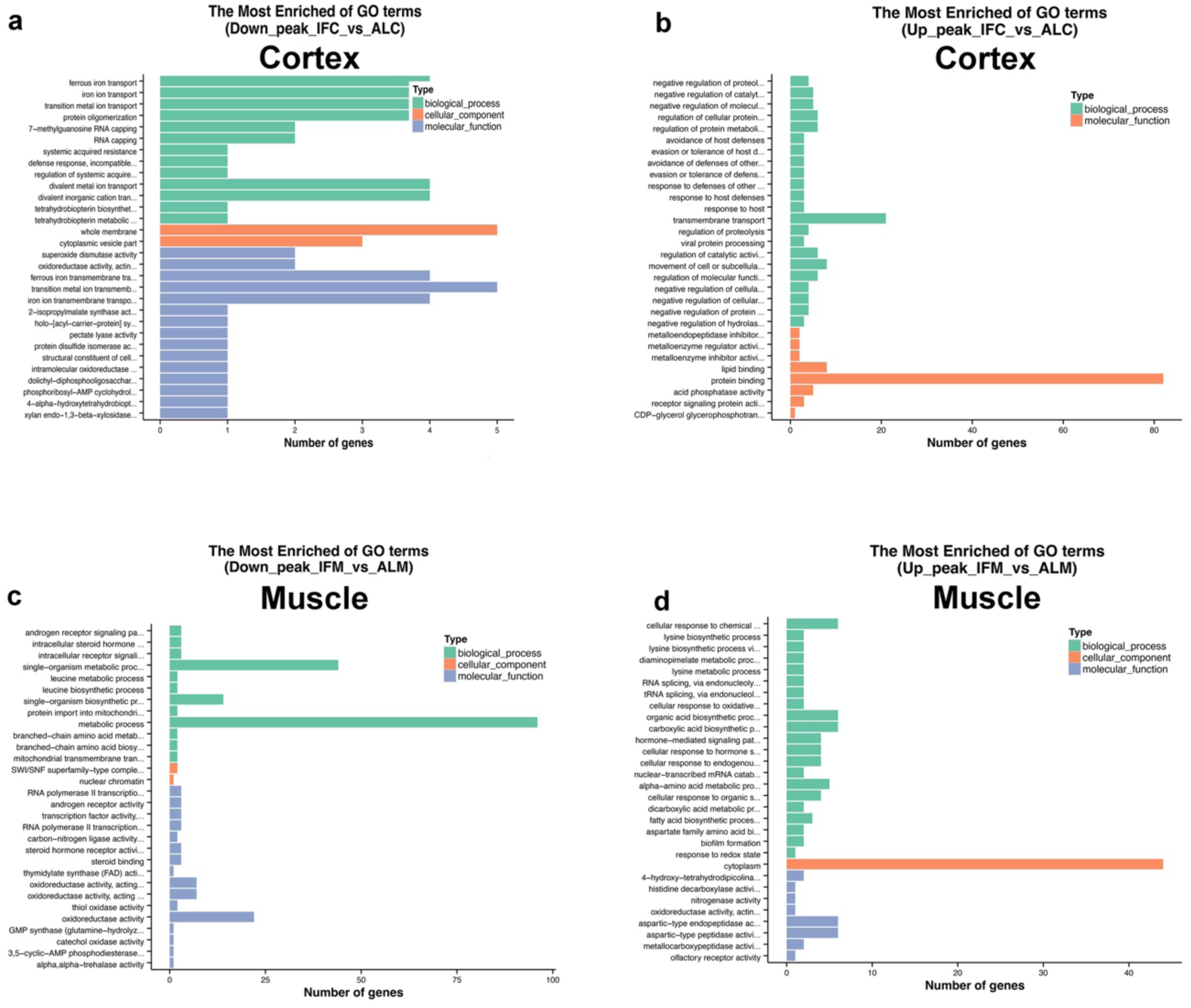
Gene Ontology (GO) Analysis of ATAC-seq. The most enriched GO terms across Biological Process (BP), Cellular Component (CC), and Molecular Function (MF) categories for each IF group compared to the AL group in cortex (**a and b**) and muscle (**c and d**). Analysis based on differentially expressed peak-related genes (DEGs) with n=8 mice/group. The height of each bar represents the number of genes, with pathways selected based on adjusted p-values (adjusted p-values < 0.05). The most enriched GO terms in BP, CC, and MF categories for each IF group compared to the AL group, based on significantly down-regulated and up-regulated peak-related genes.

**Supplementary Figure 6:**
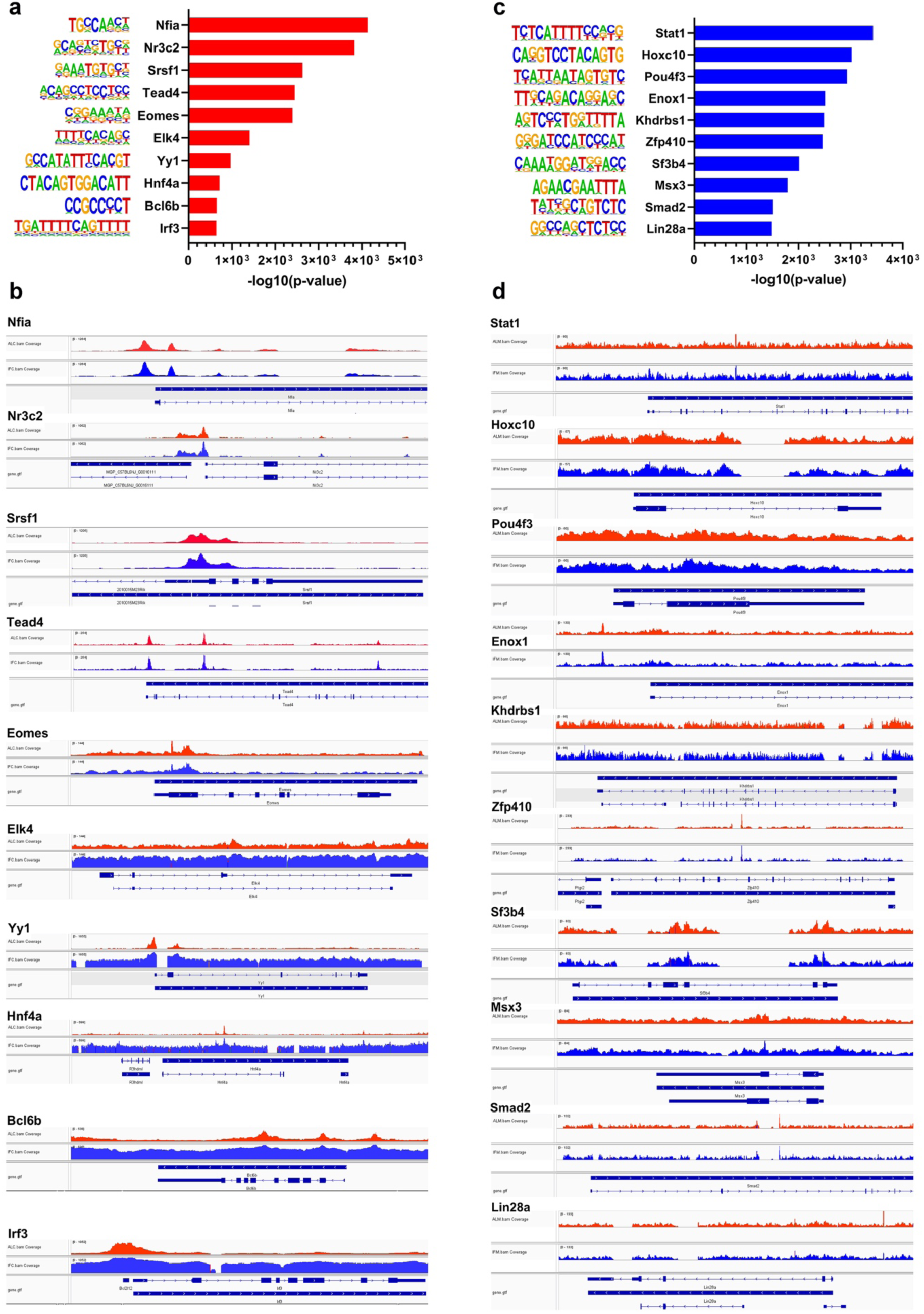
Analysis of Transcription Factor Binding Motifs in Response to Intermittent Fasting Against Baseline from HOMER Database. Top 10 most enriched transcription factors (TFs) and their corresponding motif logos in IF compared to AL in the cortex (**a**) and muscle (**c**). Transcription factors are ranked according to their -log10 p-value. Visualization of the top 10 most enriched transcription factors in IF compared to AL in the cortex (**b**) and muscle (**d**) using Integrative Genomics Viewer. Red tracks represent AL, and blue tracks represent IF. The track labeled “gene.gtf” shows the gene names, with arrows on the gene body indicating the direction of transcription. The height of the peaks represents the number of reads.

**Supplementary Figure 7:**
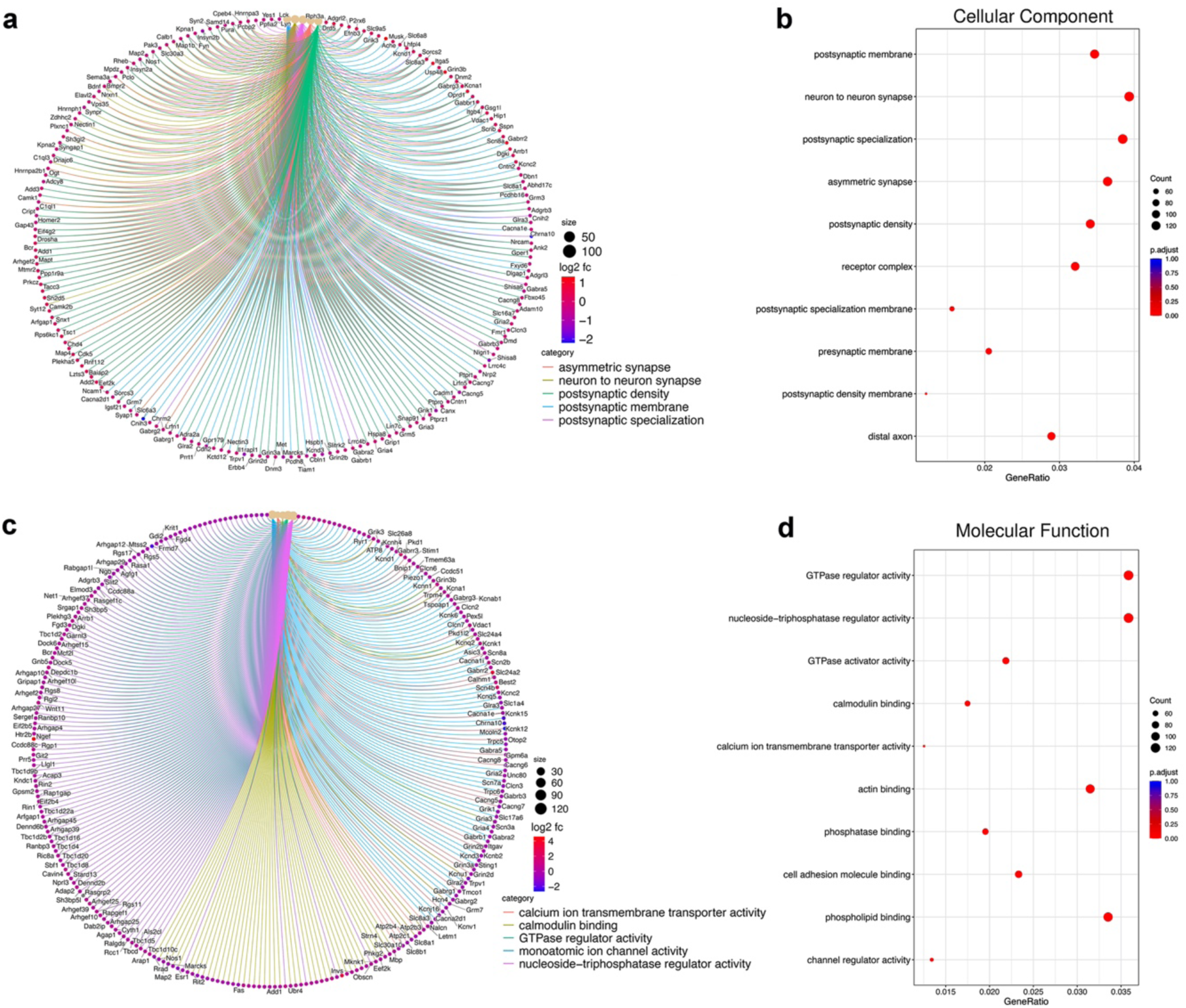
Cellular Component and Molecular Function Analysis of Genes in the Cortex from RNA-seq Data. The significantly enriched Gene Ontology (GO) terms from the categories of Cellular Component (CC) (**a and b**), and Molecular Function (MF) (**c and d**) in Circular network plot (cnetplot) for the IF group compared to the AL group in cortex (n=8 mice/group). The cnetplot highlights significantly enriched GO terms, with each node representing a gene involved in these terms. The nodes are colored according to the log2 foldchange (log2FC) of the corresponding FPKM values with colors indicating up-regulation (red) or down-regulation (blue). Terms were selected based on adjusted p-values (adjusted p < 0.05). The top 10 most enriched GO terms in the CC category (**b**) and MF category (**d**) for the IF group compared to the AL group in cortex. Terms were selected based on adjusted p-values (p.adjust < 0.05). The size of each dot represents the number of genes associated with each term, and the color indicates the adjusted p-value.

**Supplementary Figure 8:**
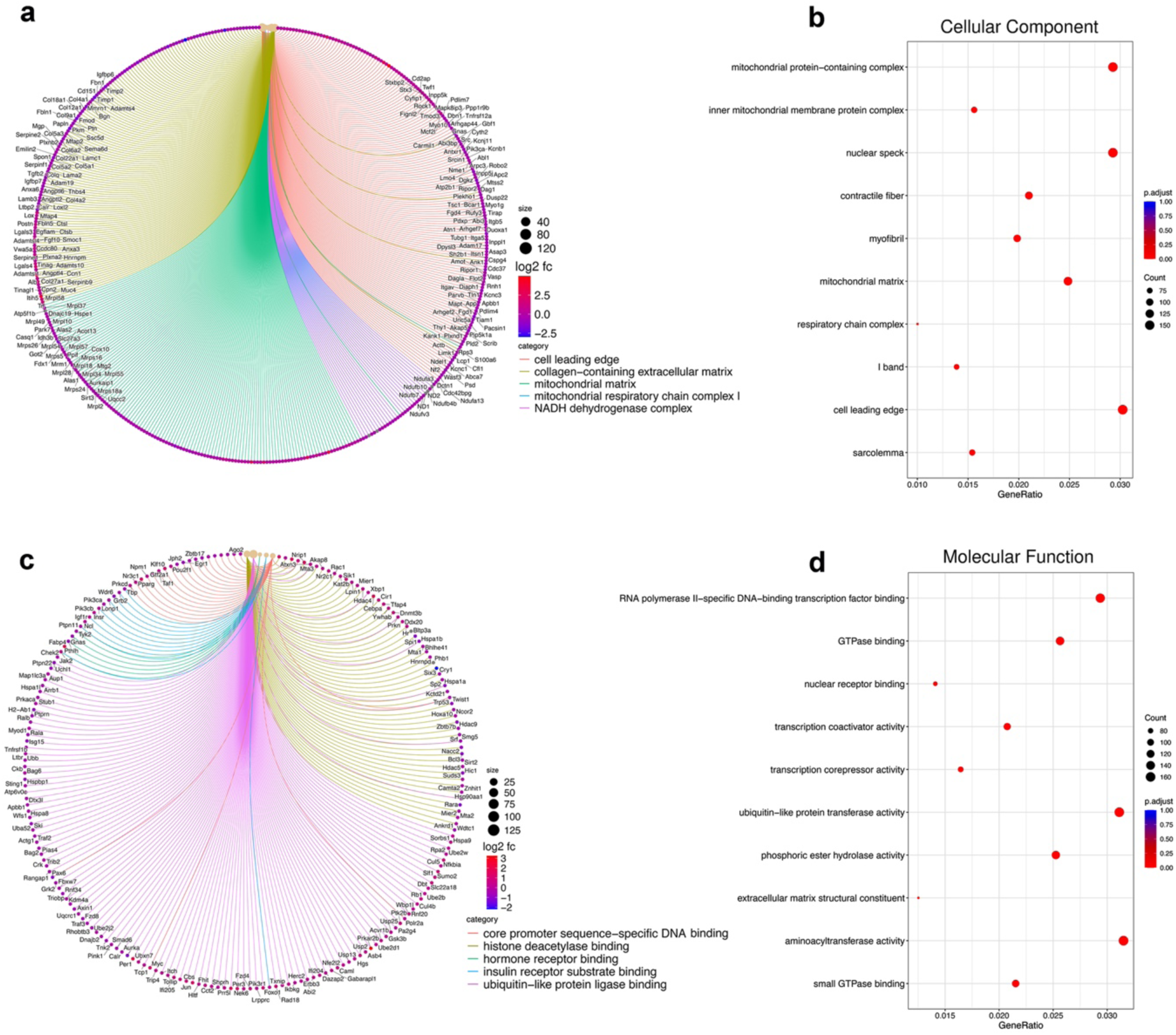
Cellular Component and Molecular Function Analysis of Genes in the Muscle from RNA-seq Data. The significantly enriched Gene Ontology (GO) terms from the categories of Cellular Component (CC) (**a and b**), and Molecular Function (MF) (**c and d**) in Circular network plot (cnetplot) for the IF group compared to the AL group in muscle (n=8 mice/group). The cnetplot highlights significantly enriched GO terms, with each node representing a gene involved in these terms. The nodes are colored according to the log2 foldchange (log2FC) of the corresponding FPKM values with colors indicating up-regulation (red) or down-regulation (blue). Terms were selected based on adjusted p-values (adjusted p < 0.05). The top 10 most enriched GO terms in the CC category (**b**) and MF category (**d**) for the IF group compared to the AL group in muscle. Terms were selected based on adjusted p-values (p.adjust < 0.05). The size of each dot represents the number of genes associated with each term, and the color indicates the adjusted p-value.

**Supplementary Figure 9:**
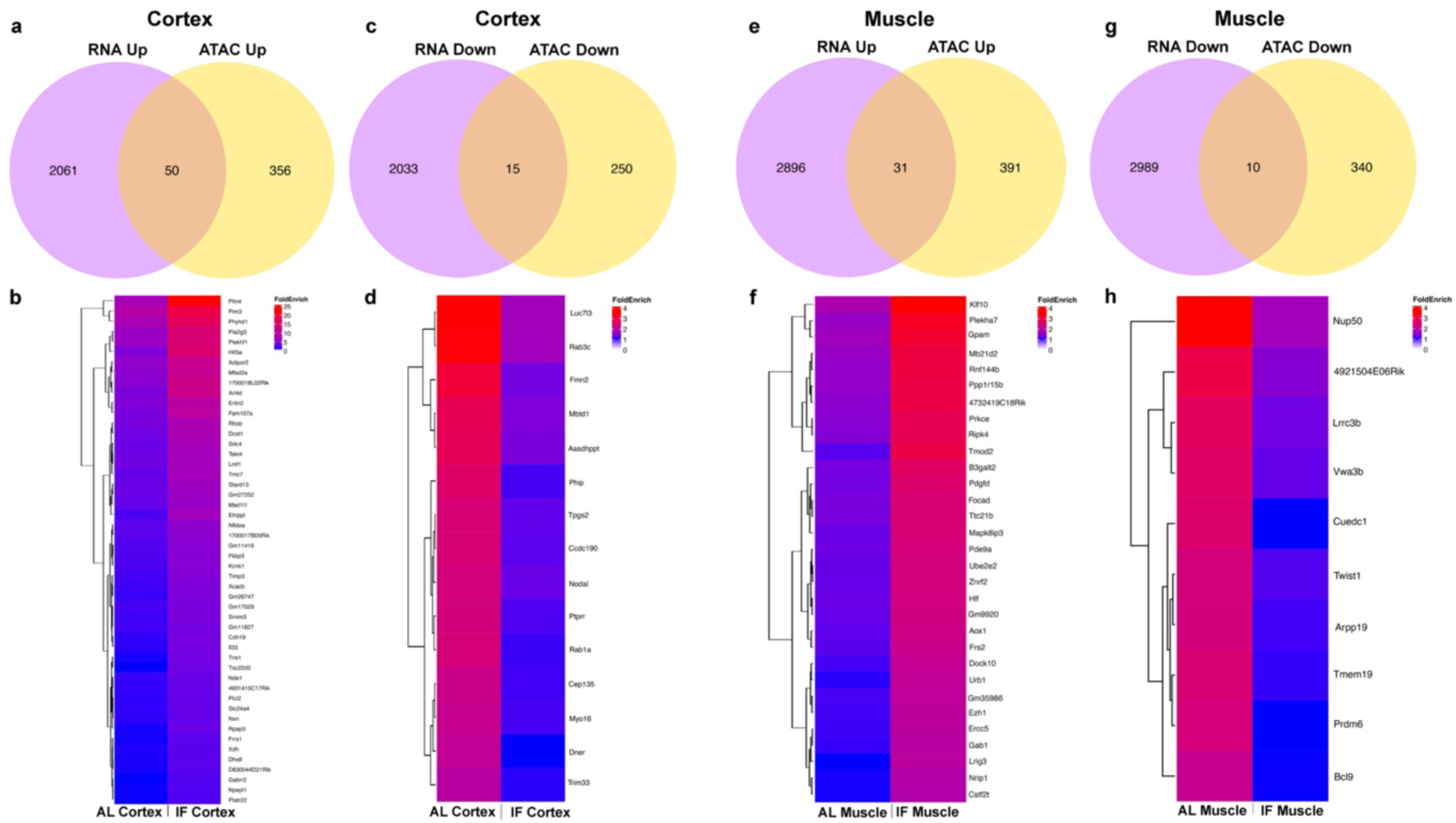
Integration of ATAC-seq and RNA-seq Results. Venn diagram of common and differential expressed genes between RNA-seq (upregulate/downregulate) and ATAC-seq (upregulate/downregulate) in both cortex (**a-d**) and muscle (**e-h**). Heatmap of common expressed genes in ATAC-seq in cortex (**b** and **d**) and the muscle (**f** and **h**). The colour scale represents fold enrichment values, with red indicating higher fold enrichment and blue indicating lower fold enrichment.

